# Microglia Rank signaling regulates GnRH function and the Hypothalamic-Pituitary-Gonadal axis

**DOI:** 10.64898/2026.02.18.706528

**Authors:** Alejandro Collado-Sole, Nozha Borjini, Jing Zhai, Gonzalo Soria-Alcaide, Cintia Folgueira, Francisco Ruiz-Pino, Celia Garcia Vilela, Victor Lopez, Yassine Zouaghi, An Jacobs, Bella Mora-Romero, Alexandra Barranco, Guillermo Yoldi, Karine Rizzoti, Guadalupe Sabio, Gema Perez-Chacon, Patricia G. Santamaria, Jose Antonio Esteban, Alberto Pascual Bravo, Rafael Fernández-Chacón, Manuel Tena-Sempere, Nelly Pitteloud, Eva Gonzalez-Suarez

## Abstract

The hypothalamic-pituitary-gonadal axis (HPG) controls pubertal development, sexual maturation, and fertility. We hereby demonstrate a new role of hypothalamic microglia in controlling the HPG axis through Rank signaling, a pathway known for its role in bone and mammary gland biology. Whole-body and microglia Rank depletion leads to hypogonadotropic hypogonadism (HH) resulting from an alteration in gonadotropin-releasing hormone (GnRH) function. In addition, we identify rare gene variants of *RANK* in patients with HH. Transcriptional profiling upon pubertal Rank loss reveals defective microglia activation and morphological alterations in the median eminence (ME), decreasing the contacts and engulfment of GnRH terminal projections and impairing GnRH responses to kisspeptin. Overall, our data uncovers the crucial role of microglia in regulating GnRH function through Rank signaling, with implications for reproductive maturation and fertility.

**One-Sentence Summary:** Microglia regulates GnRH function through RANK signaling

## Main Text

Reproductive hormones are closely related to the receptor activator of nuclear factor-kappa B (RANK) signaling pathway, regulating its activation in the bone and in the mammary gland. In the mammary epithelium, RANK ligand (RANKL) is the main paracrine mediator of progesterone, controlling mammary epithelial expansion, alveologenesis and lactation [1], [2], [3], [4]. In the bone, the drop in estrogen levels in menopause triggers overactivation of RANK signaling in osteoclasts, causing exacerbated bone turnover and osteoporosis [5]. Reproductive hormones regulate puberty onset, which is dictated by the activation of the hypothalamic-pituitary-gonadal (HPG) axis [6]. Specifically, gonadotropin-releasing hormone (GnRH) neurons are the main effectors for the control of the HPG axis in the brain, being localized in the preoptic area (PoA) of the hypothalamus, and constitute a neuronal network with projections to the median eminence (ME) [7]. Their pulsatile secretion is mainly dictated by the kisspeptin system [8] and determines the patterns of secretion of gonadotropins, follicle-stimulating hormone (FSH) and luteinizing hormone (LH), thereby controlling puberty onset, gonadal development and fertility.

Here, we provide mechanistic evidence supporting that microglia hypothalamic Rank signaling is involved in the activation of the HPG axis, with a primary action on the regulation of GnRH functionality. Through GnRH, microglia Rank signaling regulates gonadotropins, the production of sex hormones and, consequently, gonadal development and reproduction in both sexes. Importantly, inducible Rank deletion in puberty and adulthood disrupts the HPG axis in males. In parallel, we have identified different *RANK* gene variants in patients with congenital hypogonadotropic hypogonadism (CHH), a human syndrome that shares several phenotypic characteristics with full-body and microglia Rank-deficient mouse models, reinforcing the importance of Rank signaling within microglia in HPG regulation, pubertal onset, and fertility.

### Rank loss leads to severe hypogonadotropic hypogonadism in mice

While the role of Rank signaling in mammary gland development during pregnancy is well-established, its role during pubertal development, when mice undergo sexual maturation, remains unknown. To comprehend this aspect, we analyzed a constitutive Rank knockout mouse model lacking exons 4-7 (Rank^-/-^) [9]. A strong defect in mammary gland invasion was observed in the absence of Rank (Fig. S1a). Defective mammary invasion was confirmed in a new Rank knockout mouse model generated deleting *Rank* exon 1, Rank^e1-/-^ (for details, see Methods) (Fig. 1a). Rank^e1-/-^ mice display reduced body weight (Fig. S1b) and osteopetrosis (Fig. S1c), mimicking previous Rank loss-of-function models [9]. In contrast, targeted depletion of Rank to the mammary epithelium (using a Rank^Κ5Δ/Δ^ mouse model [10]), did not affect fat pad invasion (Fig. 1b). In fact, Rank^+/+^ or Rank^-/-^-derived mammary epithelial cells injected into the cleared fat pad of control mice, similarly invaded the fat pad (Fig. S1d).

**Fig. 1.**
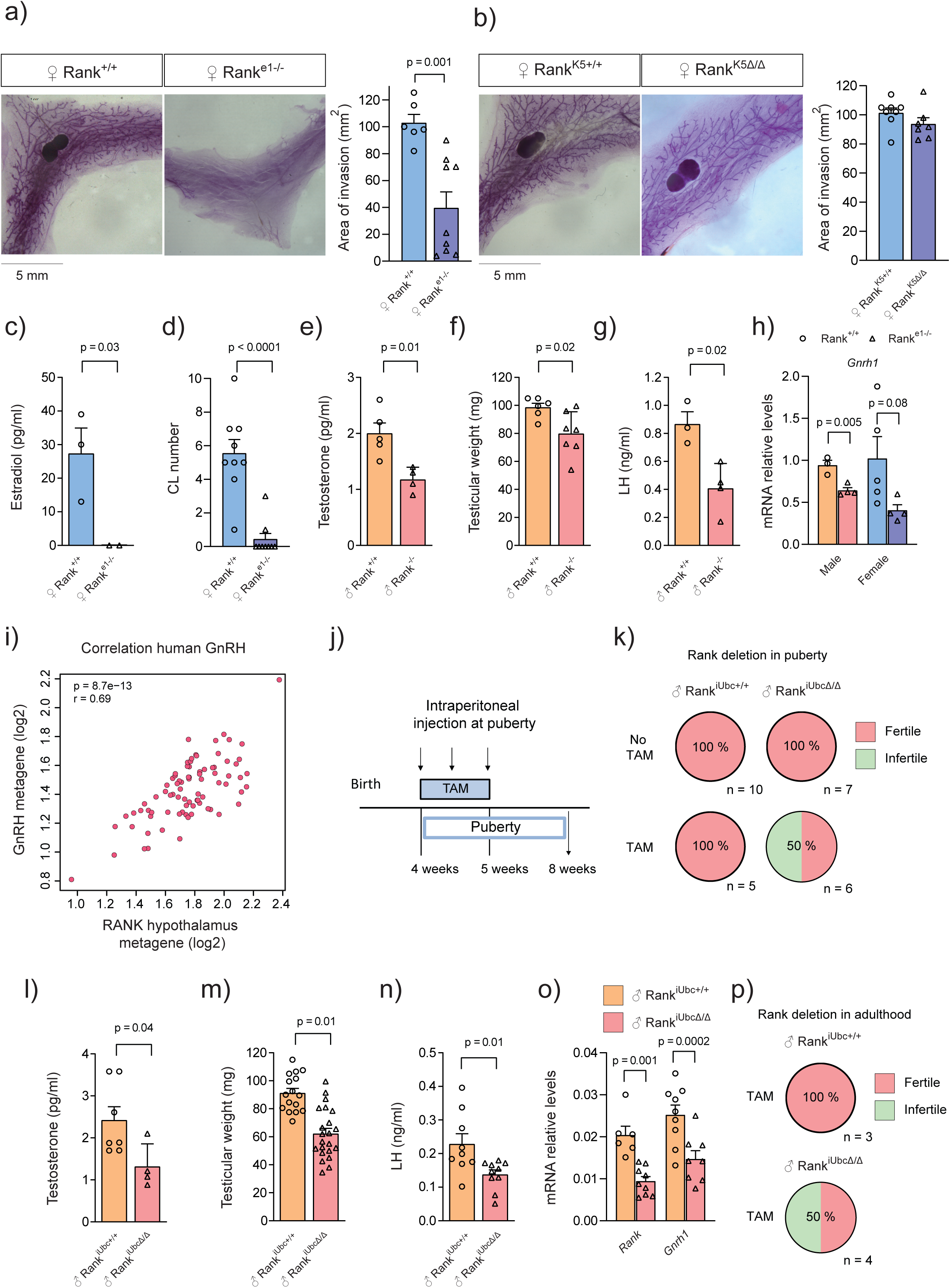
Whole-body Rank deletion induces HH in mice. (a) Representative whole mounts and quantification of epithelial invasion in mammary glands from Rank^+/+^ and Rank^e1-/-^ mice. (b) Representative whole mounts and quantification of the invaded area in mammary glands in Rank^K5+/+^ and Rank^K5Δ/Δ^ mice. (c) Estradiol (E2) levels in serum from Rank^+/+^ and Rank^e1-/-^ females. (d) Quantification of corpora lutea (CL) in Rank^+/+^ and Rank^e1-/-^ ovaries. (e) Testosterone levels in serum from Rank^+/+^ and Rank^-/-^ mice. (f) Testicular weight in Rank^+/+^ and Rank^-/-^ mice. (g) LH levels in serum from Rank^+/+^ and Rank^-/-^ males. (h) *Gnrh1* gene expression levels in the hypothalamus from Rank^ex1-/-^ male and female mice relative to control littermates. (i) Correlation between RANK metagene and GnRH metagenes analyzed in the human hypothalamus (GTEx) using GEPIA2 (Gene Expression Profiling Interactive Analysis version 2). The correlation index was calculated using Pearson correlation. (j) TAM treatment protocol to induce Rank deletion at the onset of puberty. Rank^iUbcΔ/Δ^ males and their Rank^iUbc+/+^ littermates were treated at 4 weeks of age with 3 doses of TAM (100 mg/kg) every other day for one week and euthanized 4 weeks after the first treatment. (k) Percentage of fertile Rank^iUbcΔ/Δ^ / Rank^iUbc+/+^ male mice upon Rank deletion at puberty, including TAM control and experimental treated (as shown in j) and untreated male mice. l) Testosterone levels in serum from Rank^iUbc+/+^ and Rank^iUbcΔ/Δ^ males upon Rank deletion at puberty. m) Testicular weight of Rank^iUbc+/+^ and Rank^iUbcΔ/Δ^ males upon Rank deletion at puberty. n) Circulating LH levels in blood from Rank^iUbcΔ/Δ^ males and control littermates upon Rank deletion in puberty. o) *Gnrh1* and *Rank* mRNA expression levels in the hypothalamus from Rank^iUbcΔ/Δ^ males and control littermates upon Rank deletion during puberty. p) Percentage of fertile TAM-treated Rank^iUbc+/+^ mice and Rank^iUbcΔ/Δ^ male mice upon Rank deletion in adulthood (TAM treatment as shown in Fig S3i). Analyses were performed in 8-week-old mice (a–h, k–o) and 15-week-old mice (p). Data are represented as mean +/-SEM with each dot representing a mouse; P values were calculated by unpaired two-tailed t-test and indicated when statistically significant (a-h; l-o).

As E2 is the main ovarian steroid hormone driving mammary invasion [11] and we observed an accumulation of estrogen-responsive cells in Rank^e1-/-^ mammary glands (Fig. S1e), we hypothesized that the underlying cause of the reduced mammary gland development in Rank^e1-/-^mice was related to their E2 levels [12]. Indeed, we found a drastic reduction in circulating E2 levels in Rank^e1-/-^ serum (Fig. 1c). However, cholesterol levels were similar in control and Rank^e1-/-^ serum (Fig. S1f), suggesting a defective synthesis of E2 instead of the lack of its precursor. Ovaries from Rank^e1-/-^ mice expressed lower mRNA levels of steroidogenesis proteins, responsible for E2 synthesis (Fig. S1g). This reduction was not observed in other steroidogenic tissues, such as the adrenal gland (Fig. S1h). Histological analyses showed an absence of corpora lutea in ovaries from most Rank^e1-/-^ females (up to 80% of analyzed mice) (Fig. 1d, Fig. S1i), denoting a deficiency in ovulation. Moreover, Rank-null mice had smaller (Fig. S1j) and lighter uterus than their control littermates (Fig. S1k), in line with the reduced E2 levels. Together, these results demonstrate that full-body Rank deletion results in defective ovulation and E2 synthesis. Next, we analyzed whether Rank loss would also impact gonadal development in males. Rank^-/-^ male mice displayed lower levels of testosterone in serum (Fig. 1e) and a reduced expression of key testicular steroidogenic factors (Fig. S2a). Testes from Rank^-/-^ mice were smaller (Fig. 1f) and showed a decrease in the tubular area of seminiferous tubules (Fig. S2b), indicating the presence of hypogonadism also in Rank^-/-^ males.

As both, Rank-null males and females displayed hypogonadism, we wondered whether the hormones responsible for gonadal development, FSH and LH, might be altered. Actually, circulating LH was reduced in Rank^-/-^ males (Fig. 1g) and gene expression analyses of the pituitary glands showed a reduction of approximately 60% in *Lhb* and *Fshb* mRNA levels while gene expression of other pituitary hormones such as *Prl*, *Gh* and *Tshb* was not altered (Fig. 2Sc). In Rank^e1-/-^ females, lower *Lhb, Fshb* and *Prl* mRNA levels were detected in the pituitary (Fig. S2d). The reduction of *Prl* mRNA in females, but not in males, is likely driven by the low levels of E2 (Fig. 1c), which regulates the synthesis and secretion of prolactin [13]. FSH and LH levels are tightly regulated by pulsatile secretory patterns of GnRH released from hypothalamic GnRH neurons [7] and indeed, the analysis of *Gnrh1* mRNA in Rank^e1-/-^ hypothalamus pointed to a clear downregulation of this neuropeptide in male and female mice (Fig. 1h). In line with the key role of GnRH neurons in the initiation of puberty, Rank^e1-/-^ mice display delayed balanopreputial separation and vaginal opening, as external signs of puberty onset in male and female mice, respectively (Fig. S2e-f).

We then investigated RANK signaling in the human hypothalamus and its plausible role in the control of GnRH production, interrogating available transcriptomic data from human healthy hypothalamus (GTEx, n=170) [14]. To assess RANK signaling activation status, we computed RANK metagene, which included the top 100 genes co-expressed with *RANK* mRNA in the hypothalamus (Table S1). RANK metagene strongly correlated with signatures of RANK pathway activation (r > 0.69) (Fig. S2g) and, importantly, with GnRH metagene (r = 0.69) (Fig. 1h, Table S1), whereas a lower correlation (r < 0.35) with other hypothalamic neuropeptides, such as growth-hormone and corticotropin-releasing hormones (GHRH and CRH), was found (Fig. S2h, Table S1). Analyses of gene expression in mouse hypothalamus corroborated the specific correlation of *Rank* with *Gnrh1* expression (Fig. S2i).

In sum, these results strongly suggest a control of GnRH levels by RANK signaling. As a consequence, Rank loss induces severe HH in mice, a condition characterized by defective production of sex hormones and delay in sexual maturation due to a reduced gonadotropin drive.

### Patients with congenital hypogonadotropic hypogonadism harbor rare *RANK* variants

Global Rank-deficient mice show striking physiological similarities with patients with CHH, who lack pubertal development due to defective migration of GnRH neurons or GnRH secretion. In approximately 50% of CHH cases, no specific gene mutation has been documented [15]. Therefore, the identification of the whole spectrum of molecular alterations underlying the defective regulation of GnRH neurons in CHH remains an unmet medical need. Whole-exome sequencing of 564 unrelated CHH probands identified six rare heterozygous variants in *TNFRSF11A* (RANK) in 6 unrelated probands who do not carry any known CHH gene defect, each with a minor allele frequency (MAF) < 0.1% in gnomAD v4.0 [16], collectively found in 1% of the cohort (n = 6), including four CHH individuals with anosmia (termed Kallmann syndrome) (Table 1). Two variants—p.K240E and p.E382G—met stringent in silico criteria for predicted pathogenicity (Combined Annotation Dependent Depletion (CADD) > 20 and AlphaMissense > 0.5) and mapped to highly conserved residues (Fig. S3a), suggesting functional consequences. The p.K240E variant (Proband B) segregated with delayed puberty in a sibling and was also present in an unaffected mother, consistent with incomplete penetrance and variable expressivity (Fig. S3b). Based on these findings, we expanded our analysis to rare predicted-deleterious variants in genes comprising the RANK hypothalamic metagene, a transcriptional module associated with RANK pathway activation (Fig. S2g). We identified 76 unique heterozygous rare variants predicted to be pathogenic across 73 probands (13.1%) (Table S2), including 16 putative loss-of-function events (stop, gains, frameshifts, or essential splice-site mutations). Several metagene components were recurrently affected, including *ATP8B4* (n = 7), *ENTPD4* (n = 5), *LRMP* (n = 4), *RGS3* (n = 4), *MEF2A* (n = 4), and *PTGS1* (n = 4). Two individuals carried multiple damaging variants in distinct metagene components, raising the possibility of oligogenic inheritance in a subset of CHH cases (Table S2). Notably, 70% of the probands harboring a rare variant in *RANK* or the *RANK* metagene did not carry rare variants in known CHH genes. Together, the identification of damaging variants in *RANK* and *RANK* hypothalamic metagenes in CHH patients, alone or in combination—alongside the reproductive phenotypes observed in *Rank*-deficient mouse models—supports a critical role for RANK signaling in the regulation of the hypothalamic–pituitary–gonadal (HPG) axis in humans.

**Table 1.** Clinical and genetic characteristics of CHH probands harboring rare *RANK* gene variants.

| Predicted pathogenicity | Proband | Sex | Diagnosis | *RANK* mutation | dbSNP ID | MAF | CADD | Alpha missense | Inheritance | Mutations in known CHH genes |
| --- | --- | --- | --- | --- | --- | --- | --- | --- | --- | --- |
| Tier I: Likely pathogenic | A | M | CHH reversal | c.1145A>G [p.E382G] | rs200791079 | 0.00003422 | 29.2 | 0.6398 | No information | / |
|  | B | M | KS | c.718A>G [p.K240E] | rs148185533 | 0.0007522 | 27.7 | 0.5288 | Familial | / |
| Tier II: Uncertain / likely benign | C | M | CHH | c.323C>T [p.T108I] | rs376617343 | 0.00000195 | 19.8 | 0.113 | Sporadic | / |
|  | E | M | KS | c.932C>T [p.T311I] | rs764561352 | 0.00002074 | 5.328 | 0.0945 | Familial | / |
|  | F | F | KS reversal | c.543A>T [p.R181S] | rs762733251 | 0.00002474 | 0.141 | 0.1393 | Familial | / |
|  | G | M | KS | c.766A>T [p.S256C] | . | 0 | 0.217 | 0.113 | No information | / |

Tier I (highlighted in grey) includes rare sequence variants predicted to be deleterious, with CADD scores > 20 and AlphaMissense scores > 0.5, indicating stronger pathogenic potential. Abbreviations: CADD, Combined Annotation Dependent Depletion; MAF, maximum minor allele frequency in gnomAD; HH, hypogonadotropic hypogonadism; KS, Kallmann syndrome (CHH with anosmia or hyposmia); M, male; F, female; /, absent.

### Postnatal Rank deletion suppresses the HPG axis in male mice, leading to infertility

Impaired migration of the GnRH neurons from the olfactory placode during embryogenesis is one of the causes of CHH, which is accompanied by anosmia in patients with Kallmann syndrome [17]. The fact that some CHH patients carrying *RANK* mutations are normosmic (Table 1) suggest that RANK may regulate GnRH homeostasis postnatally. To explore this hypothesis, we generated a tamoxifen (TAM) inducible loss-of-function mouse model (Rank^iUbcΔ/Δ^) that allows Rank deletion after GnRH neuron migration. Since TAM is a selective estrogen receptor modulator [18], known to affect the pituitary-ovary axis in females by itself [19], we used TAM-treated male mice in the subsequent experiments. Rank^iUbcΔ/Δ^ and their littermate controls Rank^iUbc+/+^ males were treated with TAM for one week at pubertal onset (four weeks postnatally) and analyzed at eight weeks when they had reached adulthood (Fig. 1j). Unlike constitutive Rank-null mice, Rank^iUbcΔ/Δ^ males did not display osteopetrosis (Fig. S3c), nor reduced body weight (Fig. S3d). Noticeably, TAM-treated Rank^iUbcΔ/Δ^ male were sterile or subfertile, as evidenced by their incapacity to breed with control female mice (Fig. 1k) or the longer time needed to fertilize females (Fig. S3e), in contrast to control Rank^iUbc+/+^ TAM-treated and untreated Rank^iUbcΔ/Δ^ and Rank^iUbc+/+^ male mice. Fertility was not altered in TAM-treated Rank^iUbc+/+^ mice, reflecting the minimal impact of TAM itself on reproduction (Fig. 1k, Fig. S3e). Rank depletion at puberty onset reduced testosterone levels (Fig 1l), testicular weight (Fig. 1m) and the area of seminiferous tubules (Fig. S3f), suggesting impairment of spermatogenesis. Moreover, Rank^iUbcΔ/Δ^ males showed reduced testicular levels of mRNAs of key steroidogenic factors (Fig. S3g), and *Fshb* and *Lhb* mRNA levels in the pituitary glands, compared to TAM-treated control Rank^iUbc+/+^ males (Fig. S3h), as well as lower levels of circulating LH (Fig. 1n). In contrast, no major changes in pituitary gene expression of non-gonadotropin hormones (*Prl*, *Gh*, *Tshb*) were observed (Fig. S3h). Importantly, *Gnrh1* and *Rank* mRNA levels in Rank^iUbcΔ/Δ^ mice were reduced in the hypothalamus, confirming the depletion of *Rank* in the hypothalamic tissue (Fig. 1o). Next, we investigated the consequences of depleting Rank after mice had reached sexual maturation (Fig. S3i). Likewise, adult Rank deletion led to lower mRNA of *Gnrh1* and *Rank* in hypothalamic levels (Fig. S3j) and induced hypogonadism, reflected by reduced testicular weight (Fig. S3k) and infertility in half of Rank^iUbcΔ/Δ^ mice (Fig. 1p). Together, these results corroborate that Rank signaling not only controls the HPG axis during embryogenesis, but also during puberty and adulthood, through a tight regulatory network.

### Microglia, the main source of Rank in the hypothalamus, regulates the HPG axis

Next, we assessed whether the defects in the HPG axis observed upon global Rank depletion could be due to an intrinsic role of Rank signaling in GnRH neurons. We generated Rank^Gnrh1Δ/Δ^ mice to specifically delete Rank in GnRH neurons. However, these mice did not exhibit any defects in testicular or uterus weight (Fig. S4a-b), area of seminiferous tubules in the testes (Fig. S4c) or presence of corpora lutea in ovaries (Fig. S4d), indicating that Rank loss in GnRH neurons does not impair the establishment of the HPG axis. Therefore, we asked which Rank-expressing cell population in the hypothalamus might be responsible for regulating the GnRH neurons and the reproductive axis. Analysis of scRNAseq datasets from the mouse [20] and human hypothalamus [21] revealed that the *Rank/RANK* genes and corresponding metagenes are almost exclusively expressed in the microglia (Fig. 2a, Fig. S5a). In fact, the RANK metagene exhibited enrichment in pathways related to myeloid differentiation, cytokine production and phagocytosis which are well-established pathways linked to microglia (Fig. S5b) and correlated (r = 0.95) with a gene signature comprising microglial markers from scRNAseq data [22], with nine genes overlapping between both signatures (Fig S5c-d). We then hypothesize that microglia expressing Rank regulates GnRH neurons and the establishment of the HPG axis. As there is no current evidence demonstrating a role for microglia in the regulation of GnRH neurons, we used the Csf1r inhibitor, PLX3397, which depletes microglia, and assessed its impact on the reproductive system. Control female and male mice were treated with PLX3397 for four weeks starting at puberty onset. PLX3397 treatment eliminated completely microglia (Iba1⁺ cells) in the hypothalamus in both sexes (Fig. S5e). Strikingly, microglial depletion disrupted the development of reproductive tissues, resulting in reduced testicular weight (Fig. S5f), a marked decrease in seminiferous tubule area (Fig. S5g), and a lower number of corpora lutea in the ovaries of PLX3397-treated mice (Fig. S5h). These findings support a role of microglia in the regulation of the HPG axis.

**Fig. 2.**
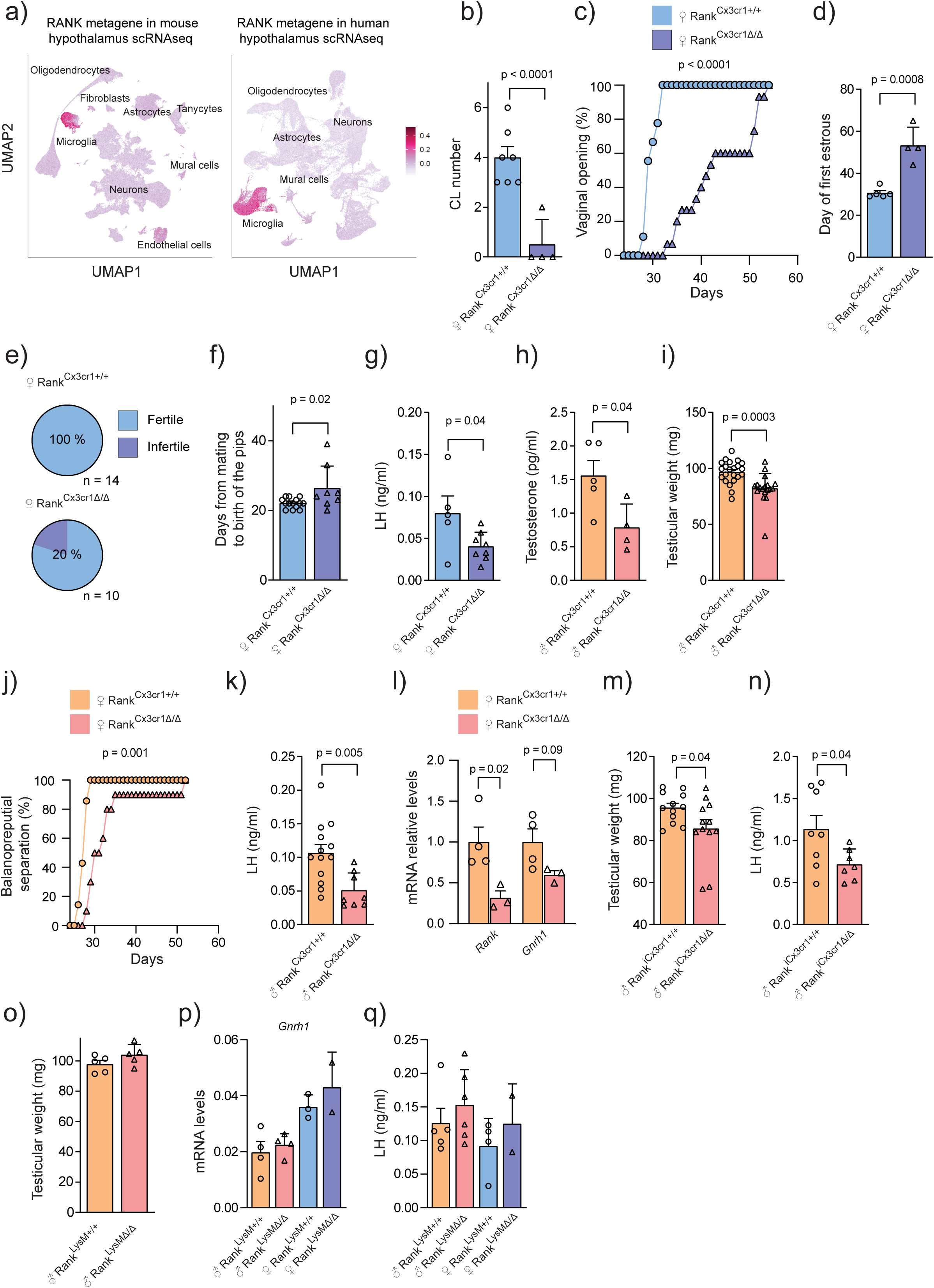
Microglia Rank loss impairs the HPG axis. (a) Left plot: *Rank* expression in mouse hypothalamus from an integrated analysis of 17 scRNAseq datasets, including embryonic and adult mice (PMID: 36266547). Right plot: *RANK* expression in human hypothalamus scRNAseq dataset from human embryos (GEO: GSE169109). (b) Quantification of corpora lutea (CL) in control and Rank^Cx3cr1Δ/Δ^ ovaries. (c) Graph showing the cumulative percentage of mice that achieved vaginal opening state. P-values calculated by the Gehan-Breslow-Wilcoxon matched-pairs test are shown (WT vs. Rank^Cx3cr1Δ/Δ^, χ² = 22.45, P < 0.0001, n = 9 and n = 13). (d) Day of the first estrous cycle determined by daily vaginal smear after vaginal opening from control and Rank^Cx3cr1Δ/Δ^ females. (e) Percentage of fertile Rank^Cx3cr1+/+^ (n = 14) and Rank^Cx3cr1Δ/Δ^ female mice. (f) Days from mating to the birth of the pups of animals from panel e excluding the infertile mice from panel e (g) Circulating LH levels in blood from Rank^Cx3cr1Δ/Δ^ and control Rank^Cx3cr1+/+^ females. (h) Testosterone levels in serum from Rank^Cx3cr1Δ/Δ^ male mice and control littermates. (i) Testicular weight of Rank^Cx3cr1Δ/Δ^ male mice and control littermates. (j) Graph showing the cumulative percentage of males that achieved balanopreputial separation in Rank^Cx3cr1Δ/Δ^ male mice and controls. Statistical differences were tested using the Gehan-Breslow-Wilcoxon matched-pairs test: Rank^Cx3cr1+/+^ vs. Rank^Cx3cr1Δ/Δ^, χ² = 7.897, P = 0.0034, n = 7 and n = 8. (k) Circulating LH levels in blood from Rank^Cx3cr1Δ/Δ^ and control Rank^Cx3cr1+/+^ males. (l) *Gnrh1* and *Rank* gene expression levels in the hypothalamus from Rank^Cx3cr1Δ/Δ^ relative to control male mice. (m) Testicular weight of Rank^iCx3cr1+/+^ and Rank^iCx3cr1Δ/Δ^ upon TAM treatment at puberty. (n) Circulating LH levels in blood from TAM-treated at puberty Rank^iCx3cr1+/+^ and Rank^iCx3cr1Δ/Δ^ mice. (o) Testicular weight in Rank^LysMΔ/Δ^ and control littermates. (p) *Gnrh1* mRNA levels of Rank^LysM+/+^ and Rank^LysMΔ/Δ^ hypothalamus of male and female mice. (q) LH levels in blood from Rank^LysM/+^ and Rank^LysMΔ/Δ^ male and female mice. Analyses were performed in 8-week-old mice (b–q). Data are represented as mean +/-SEM with each dot representing a mouse; p values were calculated by unpaired two-tailed t-test and indicated when statistically significant (b, d, f-I, k-q).

### Rank loss in microglia impairs the HPG axis in mice

To directly assess whether microglia regulate the HPG axis through Rank signaling, we generated two embryonic myeloid Rank loss-of-function mouse models: Rank^Csf1rΔ/Δ^ and Rank^Cx3cr1Δ/Δ^ (for details, see Methods). Rank^Csf1rΔ/Δ^ mice, depleted for *Rank* in myeloid progenitors, showed a similar developmental phenotype as Rank-null mice (Fig. S6a-b). Meanwhile, Rank^Cx3cr1Δ/Δ^ mice displayed no apparent phenotype in the vertebral bone (Fig. S6c), although some mice showed a deficiency in tooth eruption and decreased body weight (Fig. S6d). In both models, the loss of Rank in microglia induced HH, similar to full-body Rank deletion. Females from both models showed an almost complete absence of corpora lutea, reduced uterine weight, defective mammary gland development and increased frequency of estrogen receptor positive (ER+) cells in the mammary epithelia (Fig. 2b, Fig. S6e-i), suggesting defective E2 levels in circulation. The analysis of vaginal opening and the day of first estrous revealed a significant delay in Rank^Cx3cr1Δ/Δ^ females (Fig. 2c-d), demonstrating that microglial Rank loss interferes with the onset of puberty in females. Indeed, adult Rank^Cx3cr1Δ/Δ^ female display irregular estrous cycle with prolonged periods in diestrus (Fig. S6j). Accordingly, 20% of Rank^Cx3cr1Δ/Δ^ females were sterile (Fig. 2e), and the remaining females showed a subfertile phenotype, requiring more time to become pregnant than control females (Fig. 2f). In line with the observed dysfunctionalities of the HPG axis, Rank^Cx3cr1Δ/Δ^ females showed lower circulating LH levels (Fig. 2g) and reduced *Fshb* and *Lhb* mRNA levels in the pituitary glands (Fig. S6k). In Rank^Csf1rΔ/Δ^ females a trend toward reduced circulating LH levels was observed (Fig. S6l), together with lower levels of *Gnrh1* expression in the hypothalamus (Fig. S6m).

Similarly, Rank^Csf1rΔ/Δ^ and Rank^Cx3cr1Δ/Δ^ males displayed lower levels of circulating testosterone, decreased testicular weight and reduced area of seminiferous tubules, an indicator of impaired spermatogenesis (Fig. 2h-i and S7a-c). In fact, Rank^Cx3cr1Δ/Δ^ male mice showed a tendency to be subfertile when bred with control females (Fig. S7d-e). Rank^Csf1rΔ/Δ^ and Rank^Cx3cr1Δ/Δ^ males exhibited a delay in the balanopreputial separation (Fig. 2j, S7f), indicating an impact of microglia Rank loss on pubertal onset also in males. Consistently, Rank^Cx3cr1Δ/Δ^ male also display lower circulating LH levels (Fig. 2k) and reduced *Lhb,* but not *Fshb,* mRNA levels in the pituitary glands (Fig. S7g). Hypothalamic Rank deletion was confirmed in Rank^Cx3cr1Δ/Δ^ males and reduced *Gnrh1* expression was found in Rank^Cx3cr1Δ/Δ^ and Rank^Csf1rΔ/Δ^ males (Fig. 2l, S7h).

Moreover, to confirm the specific role of microglia Rank signaling in the postnatal control of the HPG axis, we developed two independent TAM-inducible microglia loss-of function mouse models: Rank^iCx3cr1Δ/Δ^ and Rank^iTmem119Δ/Δ^ (see Methods). Importantly, microglial Rank depletion at pubertal onset in males (as shown in Fig. 1j) did not affect body weight during pubertal growth in any of the two models (Fig. S7i-j). Rank^iCx3cr1Δ/Δ^ adult mice showed lower testicular weight (Fig. 2m) and reduced seminiferous tubule area (Fig. S7k). Indeed, a reduced number of epithelial cells within the seminiferous tubules was detected, confirming defective testicular histology (Fig. S7l). In line with this defective gonadal development, in the Rank^iCx3cr1Δ/Δ^ pituitary tissue lower circulating levels of LH (Fig. 2n) were observed, along with a reduction of *Lhb*, but not of *Fshb*, mRNA expression (Fig. S7m). A trend toward reduced testicular weight (Fig. S7n) and circulating LH levels (Fig. S7o) was also observed in Rank^iTmem119Δ/Δ^ males. Together, results from the four independent models, demonstrate that embryonic or postnatal microglia Rank loss leads to HH.

### Rank loss in peripheral myeloid cells does not impair fertility or sexual maturation

*Csf1r* and *Cx3cr1* models also display recombination in peripheral cells [23]. To assess whether Rank loss in peripheral myeloid cells might contribute to the HH phenotype, we generated Rank^LysMΔ/Δ^ mouse model, [23], which maintains *Rank* expression in the hypothalamus (Fig. S8a), but recombines efficiently in peripheral myeloid cells [23]. Rank^LysMΔ/Δ^ mice, displayed no changes in body weight and no evidence of bone defects (Fig. S8b-c). In contrast to previous microglia-specific mouse model, Rank^LysMΔ/Δ^ mice showed unaltered regulation of the HPG and were fertile (Fig. S8d-e). The day of vaginal opening (Fig. S8f), the weight of uterus or testes (Fig. S8g and Fig. 2o), and gonadal histology (Fig. S8h-i) were not affected by the loss of Rank in peripheral myeloid cells. Additionally, comparable mammary gland development during puberty and similar ER+ cell frequency within mammary glands was observed in Rank^LysMΔ/Δ^ females, both indicative of normal systemic E2 levels (Fig. S8j-k). The levels of hypothalamic *Gnrh1* mRNA levels (Fig. 2p) and circulating LH levels (Fig. 2q) in Rank^LysMΔ/Δ^ males and females were comparable to those of controls, indicative of an intact HPG regulation upon Rank loss in the peripheral myeloid cells. Since Rank expression is limited to microglia in the hypothalamus, and the HH phenotype was observed in Rank^Csf1rΔ/Δ^, Rank^Cx3cr1Δ/Δ^, Rank^iCx3cr1Δ/Δ^ and Rank^iTmem119Δ/Δ^ mice, but absent in Rank^LysMΔ/Δ^ mice (where Rank is deleted in peripheral myeloid cells), our findings demonstrate that Rank signaling in microglia regulates *Gnrh1* mRNA expression and, in turn, modulates the HPG axis and fertility.

### Transcriptomic profiling at the single-cell level reveals reduced microglia activation upon Rank loss

To elucidate the molecular consequences and pathways affected by Rank loss on the hypothalamic microglia, single-cell transcriptional profiling was performed on the hypothalamus from control and Rank^iUbcΔ/Δ^ males four weeks after pubertal Rank deletion. A pool of five hypothalamus per genotype was analysed after a Percoll centrifugation step to enrich for microglia. Following quality control filtering, 5,807 from control and 3,990 Rank^iUbcΔ/Δ^ cells were sequenced (70,000 reads/cell). Clustering analysis using Uniform Manifold Approximation and Projection (UMAP) and known hypothalamic cell types markers [20], [24], [25], [26], revealed twelve hypothalamic clusters including microglia, astrocytes, oligodendrocytes, among others (Fig. S9a-b). Remarkably, Rank^iUbcΔ/Δ^ microglia displayed the highest number of differentially expressed genes (DEGs) compared to the corresponding controls (101 DEGs, 83 downregulated p-adj < 0.05), while other hypothalamic populations, including perivascular macrophages (PVMs), or astrocytes displayed very few DEGs between genotypes, reinforcing the microglia as the most affected hypothalamic population upon Rank loss (Fig. 3a, Table S3).

**Fig. 3.**
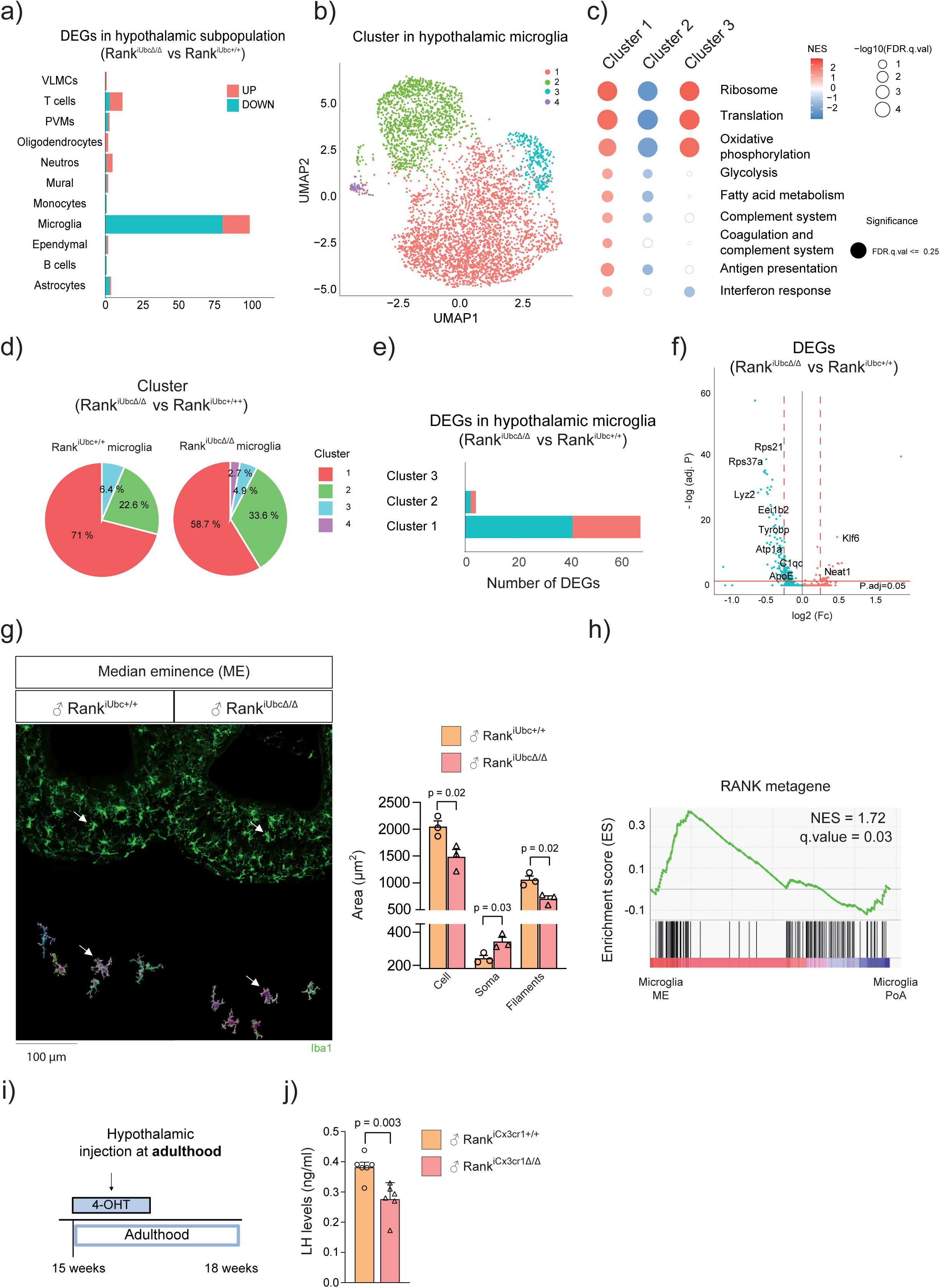
Loss of Rank induces transcriptomic changes in the microglia and alters its morphology in the ME. (a) Bar plots showing differential expression genes (DEGs) upon pubertal Rank loss between Rank^iUbcΔ/Δ^ and Rank^iUbc+/+^ male mice within each hypothalamic cluster by scRNAseq. VLMCs = Vascular and leptomeningeal cells, PVMs = Perivascular macrophages. (b) UMAP of hypothalamic microglia showing four different subpopulations by the combination of Ranki^UbcΔ/Δ^ and Rank^iUbc+/+^ cells upon Rank deletion in puberty. The fourth microglia cluster did not pass the quality control due to its limited abundance (57 out of 2,100 microglia cells) and the expression of the female-specific gene *Xist* in males (c) Bubble plot depicting the variation of selected gene set signatures from cluster 1, 2 and 3. Filled bubbles indicate enriched significant gene signatures (FDR < 0.25). Red and blue colors indicate upregulated (NES>0) and downregulated (NES<0). (d) Circle charts of all microglia clusters by percentage in Rank^iUbcΔ/Δ^ and control mice. (e) Bar plot showing differential expression genes (DEGs) between Rank^iUbcΔ/Δ^ versus Rank^iUbc+/+^ mice in each microglia cluster. f) Volcano plots of DEGs in Rank^iUbcΔ/Δ^ versus Rank^iUbc+/+^ in microglia. Significant DEGs above the horizontal red line based on P-adjusted value < 0.05, blue and red bubbles represent significantly downregulated or upregulated genes, respectively. Fc, fold-change.. (g) Representative immunofluorescence (IF) images of hypothalamic microglia cells (Iba1+ in green) in the median eminence (ME) from Rank^iUbcΔ/Δ^ and Rank^iUbc+/+^ male mice upon pubertal Rank loss, Reconstructed microglia images generated using IMARIS software (marked with white arrow) and quantification of microglial morphological parameters including total microglial area, cell body area and microglial filament area are shown. (h) GSEA of the RANK hypothalamic metagene between ME and preoptic area (PoA) microglia. (i) TAM treatment procedure to delete Rank in the ME. Mice (12-15 weeks) were treated at adulthood with 10 mM 4-hydroxytamoxifen (4-OHT) intracranially in the ME and were euthanized 4 weeks after the first treatment. (j) Circulating LH levels in blood from Rank^iCx3cr1Δ/Δ^ and control littermates after 4 weeks of intracranial injection with 4-OHT. Data are represented as mean ± SEM, with each dot representing a mouse, P values were calculated using unpaired two-tailed t-test and indicated when statistically significant (g, j). Rank^iUbcΔ/Δ^ and Rank^iUbc+/+^ male mice were TAM treated as shown in Fig. 1j (a-g)

Focusing on the microglia cluster, three main subpopulations of hypothalamic microglia were found (Fig. 3b). Cluster 1 represents the predominant population enriched in common microglial homeostatic genes, such as *P2ry12* and *Cx3cr1*, as well as phagocytic genes, including *Tyrobp, C1qa and Lys2* (Fig. S9c). Gene set enrichment analysis (GSEA) indicated an active population with higher protein synthesis capacity, metabolism, antigen presentation, interferon response and complement system processes (Fig. 3c, Table S4). Indeed, cluster 1 shares transcriptional similarities with metabolic and proliferative microglia found in embryogenesis, during inflammatory/injury response and in aging [27], [28] (Fig. S9d, Table S4). Cluster 2 displays downregulation of most pathways analyzed, including translation and metabolism, and negative associations with proliferative and inflammatory pathways (Fig. 3c, Fig. S9d, Table S4). Cluster 2 is characterized by the expression of the transcription factor, *Mef2a*, a regulator of microglia development and synapses [29], and the microglia-specific receptor *Tgfbr1* [30] (Fig. S9c). Cluster 2 displays a “mirror” image of cluster 1 (Fig. 3c, Fig. S9d, Table S4), denoting an “inert/inactive” hypothalamic-specific microglia subcluster. Cluster 3 represents metabolically active microglia, similar to cluster 1, but with less inflammation and a notable enrichment in ribosomal and respiratory pathways (Fig. 3c, Fig. S9d, Table S4).

Comparison of the relative frequencies of microglial clusters between genotypes revealed that Rank loss leads to a reduction in the active cluster 1 (from 71% to 58%), accompanied by an increase in the “inactive” cluster 2 abundance (from 22.6% in controls to 33.6% in Rank^iUbcΔ/Δ^ mice) (Fig. 3d). Moreover, the majority of DEGs were found downregulated in the whole microglia as well as in cluster 1 (Fig. 3a, e-f, Table S3, Table S5) supporting the role of Rank signaling in maintaining active hypothalamic microglia. Rank^iUbcΔ/Δ^ microglia showed downregulation of key mediators of the Trem2/Tyrobp signaling pathway (*Tyrobp, ApoE*), the complement system (*C1qa, C1qc)*, phagocytic system (*Lyz2, Gpr84, S100a8, S100a9, Ctss*), and a reduction of mediators of protein translation (*Eef1b2, Eef1d)*, mitochondrial respiration and neurodegenerative disorders (Fig. 3f, Fig. S9e, Table S3). Similar changes were observed specifically within cluster 1 upon Rank loss, where several mediators of phagocytosis, the complement system, translation and metabolism are modulated (Fig. S9f, Table S5). Together, the reduction in the relative frequency of cluster 1 and specific changes within this cluster suggest a general reduction in microglia activation upon pubertal Rank loss. Rankl stimulation in primary microglial cultures from control mice, as well as in the BV2 microglial cell line, led to the upregulation of established RANK/NF-κB target genes such as *Birc3* and *Nfkb2* as well as key mediators of the Trem2/Tyrobp phagocytic signaling pathway, including *C1qc*, *ApoE* and *LysM*—genes (Fig. S9h), reinforcing Rank signaling as a regulator of microglia activation. Altogether, these results reveal three distinct states of hypothalamic microglia and expose the central role of Rank signaling in the maintenance of homeostatic/inflammatory microglia with higher metabolic and phagocytic activity.

### Rank loss induces morphological alterations in microglia of the median eminence

Transcriptomic analyses revealed the central role of Rank signaling in the maintenance of microglia activation. Then, we assessed whether and how Rank loss affects the number and morphology of microglia in different hypothalamic regions where GnRH neurons reside. No changes in the number of microglia cells (Iba1+) were found in Rank^iUbcΔ/Δ^ mice in the PoA, where most GnRH cell bodies are located (Fig. S10a). Besides, microglia morphology in the PoA such as total microglia area, area of the cell body and area of the microglia filaments are unaffected by Rank loss (Fig. S10b), suggesting a minimal impact of Rank loss in the microglia of the PoA. Next, we evaluated microglia in the mediobasal hypothalamus, where the GnRH neuron terminals are located and secrete their ligands into the ME. We focused on the ME and on the arcuate nucleus (ARC), as Kiss1 neurons in the ARC regulate the pulsatile secretion of GnRH neurons. Although no changes in microglia abundance were observed in the ARC of Rank^iUbcΔ/Δ^ mice, an increase in the number of microglia was found in the ME (Fig. S10c). Strikingly, Rank-depleted microglia in the ME, but not in the ARC, exhibited a smaller area, bigger soma, and less processes (Fig. 3g, Fig. S10d-e). Despite the amoeboid morphology—which typically indicates microglial activation—transcriptomic analyses revealed that microglia are not functionally inflamed (Fig S9d, Table S3, Table S5). Indeed, analysis of astrocyte area in the ME revealed no differences between control and Rank-depleted mice discarding astroglial activation or inflammation (Fig. S10f).

To confirm that morphological changes are induced by Rank loss in microglia, we analyzed the microglial morphology of the ME in Rank^iCx3cr1Δ/Δ^ mice crossed with the tdTomato reporter mouse line upon TAM treatment at puberty. High recombination efficiency (tdtomato+/Iba1+) was observed in the ME of Rank^iCx3cr1Δ/Δ^ mice (Fig. S11a). Similar to the phenotype seen with ubiquitous Rank deletion, these mice displayed smaller microglia in the ME with fewer processes and larger soma (Fig. S11b), indicating that microglia Rank loss induces morphological changes in the ME microglia, consistent with alterations in microglia activation [31].

We hypothesized that Rank signaling may be more active in the ME microglia and therefore particularly sensitive to Rank loss. Indeed, we integrated two public available scRNA-seq datasets: one from the PoA [32] and another from the ME [33]. UMAP analysis of the combined microglial populations from both datasets revealed a clear distinction between ME and PoA microglia (Fig. S11c). Although Rank is expressed in microglia from both regions (Fig. S11d), ME microglia exhibited enrichment of the RANK metagene signature, indicating increased RANK pathway activity in this area (Fig. 3h). To demonstrate the importance of microglia Rank signaling in this region, we injected 4-hydroxytamoxifen (4-OHT) into the ME of adult Rank^iCx3cr1Δ/Δ^ and control mice crossed with the tdtomato reporter line (Fig. 3i). The presence of recombined microglia cells in the ME (Iba+/tdtomato+) of Rank^iUbcΔ/Δ^ confirmed the depletion of Rank in this hypothalamic population (Fig. S11e). Importantly, a reduction of circulating levels of LH in Rank^iCx3cr1Δ/Δ^ was observed (Fig. 3j), indicating a disruption in the HPG axis. Together, morphological assessments reveal a central role of Rank signaling in the regulation of microglia activation, particularly in the ME a hypothalamic area crucial in the control of the reproductive axis.

### Microglial Rank loss impairs GnRH function via reduced microglia-GnRH neuron interaction in the ME

We established that Rank loss induces specific microglial morphological changes in the ME, accompanied by a transcriptomic profile indicative of less active microglia. We then aimed to determine whether and how GnRH neurons are affected. First, we assessed the impact of embryonic Rank loss in the microglia on GnRH numbers. We performed whole-brain tissue clearing and immunostaining against GnRH using iDISCO (immunolabeling-enabled three-dimensional imaging of solvent-cleared organs) in Rank^Cx3cr1Δ/Δ^ male, where Rank was genetically eliminated in microglia from embryogenesis. We did not observe any apparent differences in the number of GnRH neurons in the whole brain (Fig. S12a). Quantification of GnRH neurons in the PoA, where most GnRH neurons reside, from independent Rank^Cx3cr1Δ/Δ^ male mice confirmed these results (Fig. S12b). Since no changes in GnRH neuron numbers were observed in the adult Rank^Cx3cr1Δ/Δ^ brain, we concluded that embryonic Rank deletion in microglia does not affect GnRH neuronal migration. Accordingly, no differences in sniffing capacity were observed in Rank^Cx3cr1Δ/Δ^ mice in habituation-dishabituation test (Fig. S12c) - a phenotype typically seen in mice with CHH and deficiencies in GnRH neuronal migration. As expected, in adult Rank^iUbcΔ/Δ^ mice, GnRH neuronal numbers in the PoA were not altered by pubertal Rank deletion (Fig. S12d), and no differences in odor tests were observed (Fig. S12e). These results highlight that Rank deletion, whether inducible or embryonic, does not alter the number of GnRH neurons nor affect their migration. Instead, the regulatory role of Rank appears to be directly related to GnRH functionality.

Next, we evaluated how the less active microglia observed upon Rank loss might communicate with GnRH neurons interfering with their function. In males, ARC Kiss1 neurons communicate with GnRH neurons at the level of their terminals near the ME [34]. Since microglia in the ME are affected by Rank loss (Fig 3g, Fig S11b), we investigated their interaction with the axon terminals of GnRH neurons. Interestingly, we observed a significant reduction in microglia–GnRH terminal contacts in the ME of Rank^iUbcΔ/Δ^ mice (Fig. 4a). Additionally, microglia in Rank-deficient mice showed reduced engulfment of GnRH content, suggesting decreased remodeling of GnRH synaptic terminals (Fig. 4a), likely due to diminished microglial activity following Rank loss as we observed in our scRNAseq analyses. Analysis of the phagocytic marker CD68 revealed fewer phagosomes within microglia, indicating reduced phagocytosis (Fig 4b), and in line with scRNAseq results (Fig. 3f). Fewer GnRH–microglia contacts in the ME and reduced engulfment of GnRH within microglia were also observed in microglia-specific Rank-deficient mice (Rank^iCx3cr1Δ/Δ^), highlighting that this phenotype results from the loss of Rank specifically in microglia (Fig. S13).

**Fig. 4.**
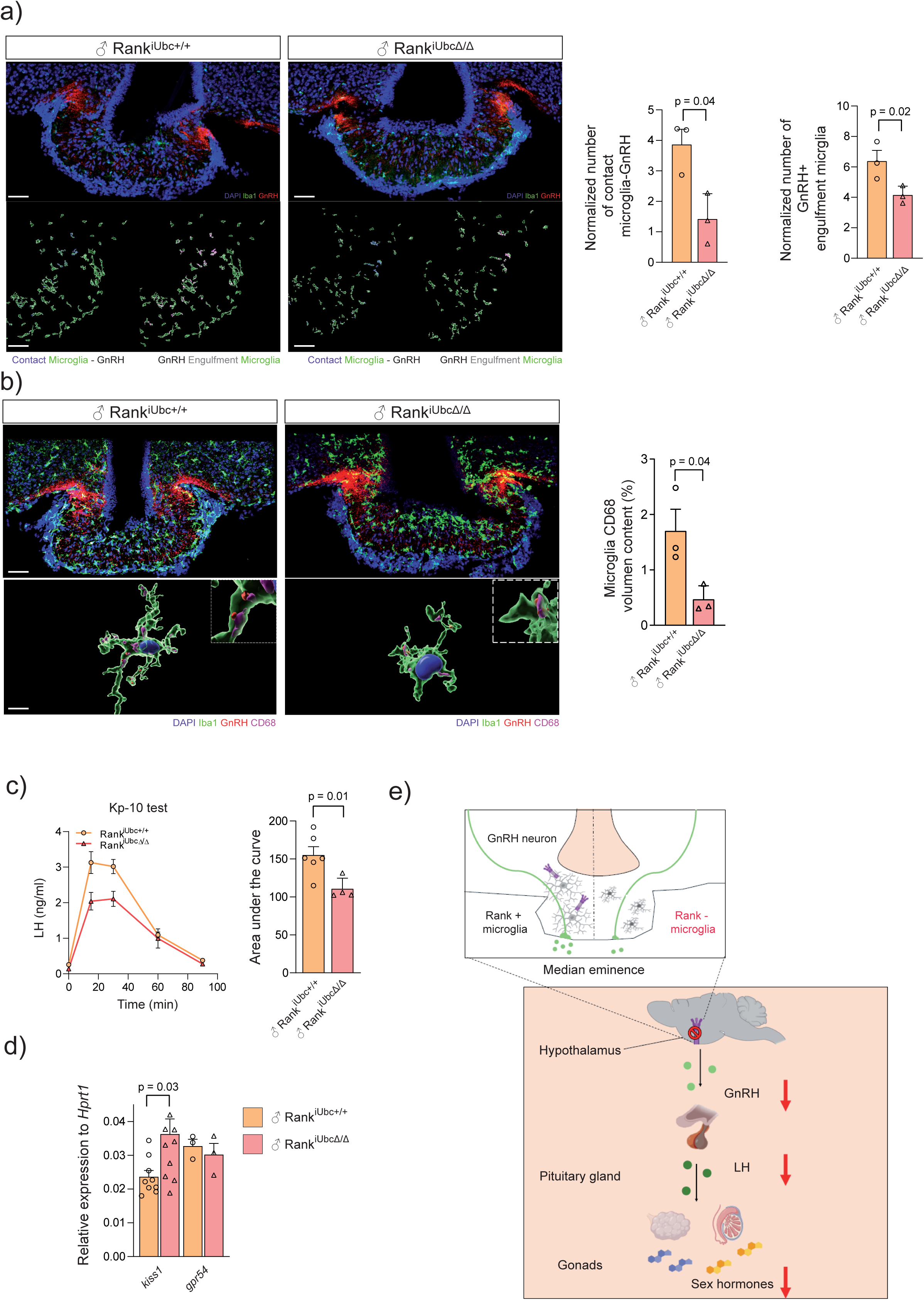
Rank loss reduces GnRH-microglia contacts and phagocytosis and disrupts GnRH function. (a) Top: Representative immunofluorescence (IF) images of microglial cells (Iba1+, green) and GnRH neurons (GnRH, red) in the median eminence (ME) of control and Rank^iUbcΔ/Δ^ mice upon pubertal Rank loss. Nuclei are labeled with DAPI (blue). Bottom: Reconstruction of microglial cells using IMARIS in the ME of control and Rank^iUbcΔ/Δ^ mice. Colocalization (left, blue) indicates areas of interaction between microglia and GnRH neurons and. Colocalization (right, pink) indicates engulfment of GnRH by microglia. Bar blots show quantification of GnRH-microglia interactions and GnRH engulfment by microglia normalized to microglial cell area. (b) Top: Representative IF images of microglial cells (Iba1+, green) GnRH neurons (GnRH, red) and CD68 (pink) in the ME of control and Rank^iUbcΔ/Δ^ mice. Nuclei are labeled with DAPI (blue). Bottom: Reconstruction of microglial cells using IMARIS in the ME of control and Rank^iUbcΔ/Δ^ mice with GnRH content (red) and CD68 (pink). Quantification of percentage of CD68 within microglia volume is shown. (c) Time-course of LH response to Kisspeptin-10 (Kp-10) in Rank^iUbc+/+^ (n = 6) Rank^iUbcΔ/Δ^ (n = 3) male mice upon Rank deletion at puberty. LH secretion was monitored at baseline (pre-injection) and at 15-, 30-, 60-, and 90-min post-injection following an acute intraperitoneal injection of Kp-10 (7.5 nmol/20 g animal). The right panel shows the quantification (AUC) from the graph on the left panel. (d) Gene expression of *Kiss1* and kisspeptin receptor (*gpr54*) in control and Rank^iUbcΔ/Δ^ hypothalamus upon Rank deletion at puberty. (e) Graphical abstract: Microglia Rank loss alters microglia activation and morphology in the median eminence, reducing GnRH–microglia contacts and impairing GnRH functionality. This, in turn, disrupts the function of the pituitary–gonadal axis. Some pictures are obtained from Biorender with modification. Data are represented as mean ± SEM, with each dot representing a mouse, P values were calculated using unpaired two-tailed t-test and indicated when statistically significant (a-d). Rank^iUbcΔ/Δ^ and Rank^iUbc+/+^ male mice were TAM treated as shown in Fig. 1j (a-d)

The function of GnRH neurons is to secrete GnRH in a pulsatile way GnRH into the portal system connecting with the pituitary gland. The main regulator of the pulsatile secretion of GnRH neurons are the subset of Kiss1 neurons from the ARC, near the ME. Therefore, we examined whether and how the GnRH/kisspeptin system was affected upon Rank loss in microglia cells. We subjected Rank^Cx3cr1Δ/Δ^, Rank^iUbcΔ/Δ^ male mice and corresponding controls to treatment with exogenous Kisspeptin-10 (Kp-10) [35] and GnRH treatment and analyzed the responsiveness of the pituitary gland *in vivo*. These pharmacological tests showed a preserved pituitary responsiveness in Rank^iUbcΔ/Δ^ /Rank^Cx3cr1Δ/Δ^ mice to exogenous GnRH (Fig. S14a-b). In contrast, a defect in the response to Kp-10 was observed in both Rank^iUbcΔ/Δ^/Rank^Cx3cr1Δ/Δ^ mice (Fig. 4a and Fig. S14c). Notably, in Rank^iUbcΔ/Δ^ hypothalamus, *Kiss1* mRNA expression was upregulated while expression of its receptor, *gpr54*, was not altered (Fig. 4b). We validated the increase of *Kiss1* mRNA in the hypothalamus of myeloid Rank-depleted mice Rank^Csf1rΔ/Δ^ (Fig. S14d). Thus, the defective response to Kp-10 observed upon microglia Rank loss is likely due to impaired GnRH neuronal secretion rather than decreased receptor expression, leading to an increase in *Kiss1* levels as a compensatory mechanism.

Together these findings indicate that Rank loss induces both morphological and functional alterations in ME microglia, leading to less phagocytic microglia and reduced contact with GnRH terminals. This impaired microglia–GnRH interaction disrupts GnRH neuron function and responsiveness to Kiss1 signaling, ultimately resulting in hypogonadism through dysfunction of the HPG axis (Fig. 4e).

## DISCUSSION

Our current data from ten independent Rank-deficient mouse models, human hypothalamic samples, patients with CHH, scRNAseq and morphological analyses of hypothalamic microglia provide strong evidence that microglia Rank signaling is a central regulator of sexual maturation and fertility through the control of hypothalamic GnRH function. Using mouse models, displaying either full-body or myeloid/microglia-restricted Rank depletion, we demonstrate that microglia Rank loss impairs microglia activation, alters microglia morphology in the ME and its interactions with GnRH neurons, disrupting GnRH functionality and leading to HH. Of note, constitutive full and myeloid/microglia Rank knockout mice exhibited bone defects and reduced body weight that might influence the hypogonadal phenotype [36]. To avoid such confounding effects, we developed inducible model allowing timed deletion of Rank at puberty onset in full-body (Rank^iUbcΔ/Δ^) or microglia (Rank^iC3xcr1Δ/Δ^ and Rank^iTmem119Δ/Δ^), which also resulted in a defective regulation of the HPG axis, without impacting body weight nor bone homeostasis. Compelling evidence, particularly in females, has documented that the influence of reduced body weight on reproduction is mainly due to changes in different neuroendocrine signals, including low leptin levels in circulation and reduced *Kiss1* expression in the hypothalamus [37]. Thus, while reduced body weight in global or microglia Rank-null mice might contribute to the suppression of the HPG axis *per se*, the fact that downregulation of GnRH also occurred in Rank^iUbcΔ/Δ^ mice or Rank^iCx3cr1Δ/Δ^, which did not show reduced body weight, argues against the possibility that the impairment of the HPG axis due to Rank ablation is merely secondary. Moreover, the absence of alterations in *Gh/Tshb/Prl* expression in Rank-null pituitary glands suggests that the reproductive phenotype is not caused by a disruption in the growth axis or other neuroendocrine axes, although a more comprehensive analyses of hormonal levels needs to be performed.

Importantly, we unveil that Rank signaling regulates GnRH within the hypothalamus and, consequently, the pituitary-gonadal axis, not only during embryogenesis and puberty causing developmental effects, but also in adulthood, indicating that Rank signaling regulates fertility even when sexual development has already been reached. Previous studies have shown that Rank signaling is essential for testicular development [38]; however, its role in the regulation of GnRH had not been specified. In women, genome-wide association studies (GWAS) have revealed single nucleotide polymorphisms (SNPs) in *RANK* and *RANKL* genes, which have been linked to a delay in the onset of the first menstrual cycle as well as menopause [39], [40], [41]. These data support a functional role for human RANK signaling throughout the female reproductive cycle.

In the embryonic models, Rank-deficient females appeared to exhibit more profound HH than males, showing significant delays in pubertal onset, infertility, and absence of ovulation, whereas males retain sperm production and are fertile. This may be attributed to sex-specific differences in the regulation of GnRH neurons or in microglial function, which are known to differ between males and females [42], [43], highlighting the need for a more thorough study of mechanism of GnRH dysfunctionality in females upon microglia Rank loss. One of the limitations of the inducible loss-of-function mouse model is its dependency on TAM treatment. In females, TAM impairs the HPG axis even at low doses [44], which prevents us from properly assessing fertility phenotypes caused by postnatal Rank loss in females. TAM, also affects the male HPG axis and at high doses induces defective spermatogenesis and hypogonadism [45], [46]. While we acknowledge that TAM treatment may affect our system, the fact that control mice treated during puberty remain fertile and do not display hypogonadism suggests that TAM has only a minimal impact on the HPG axis under our treatment protocol (low dose and washout period of recovery). Inducible, microglia specific models non-dependent on TAM would be required for a thorough study of mechanism of GnRH dysfunctionality in females upon microglia Rank loss [47].

HH is also promoted by mutations in the G protein-coupled receptor 54 (GPR54) that is expressed in GnRH neurons [48], [49]. Kisspeptins, endogenous peptide ligands of GPR54, are secreted by Kiss1 neurons and constitute the main activators of GnRH secretion. Our data demonstrate that Rank controls the function of GnRH neurons and not Kiss1 neurons, as a defective primary response of GnRH neurons to induce secretion of LH was observed under pharmacological treatment with Kp-10, as the most potent GnRH secretagogue known to date. Indeed, higher levels of *Kiss1* expression were observed in the hypothalamus upon global or myeloid Rank loss. The latter seemingly represents a compensatory mechanism to overcome the suppression of GnRH secretion, which is likely driven by the elimination of the negative feedback of sex steroids hormones upon microglial Rank ablation. In fact, androgens and estrogens have been shown to suppress *Kiss1* expression in the mediobasal hypothalamus [50], [51]; hence, HH in Rank-depleted mice is expected to enhance *Kiss1* expression. Yet, in the absence of Rank, GnRH expression is persistently low despite the compensatory rise of kisspeptin drive. To our knowledge, this is the first evidence connecting Rank signaling pathway to the central control of the HPG axis, via regulation of GnRH, thus unveiling new therapeutic targets for syndromes bound to infertility.

CHH is a rare disorder that results from the failure of pulsatile GnRH secretion, either due to defective migration of GnRH neurons from the olfactory placode during embryogenesis (i.e., Kallman syndrome), or perturbation of GnRH neuronal homeostasis at the hypothalamic site. In both cases, CHH is linked to delayed/absent puberty and infertility [17]. CHH is characterized by vast genetic heterogeneity, in which mutations in more than 30 genes, acting alone or in combination, have been described [52]. We have now identified gene variants in *RANK* and in the genes comprising the *RANK* metagene in CHH patients. This is in line with our findings in mouse models, where embryonic, pubertal and adult Rank loss leads to HH. Although we did not perform functional studies on *RANK* mutations, two of these mutations (p.K240E, p.E382G) were predicted to be damaging, showing a high CADD and AlphaMissense score, further supporting their involvement in RANK protein function. The variable expressivity and incomplete penetrance in one family carrying *RANK* mutations are consistent with a complex interaction between RANK pathway and the network of genes regulating the HPG axis.

Moreover, our results expose an intriguing link between Rank signaling and microglia functionality, as well as the involvement of microglia in the regulation of GnRH. Although the main regulation of GnRH is attributed to excitatory and inhibitory synaptic communication, such as Kiss1 system [53], growing evidence suggests that glial cells, mainly tanycytes and astrocytes, could actively contribute to this regulation [54], [55], [56]. Despite an association between hypothalamic microglia and GnRH neurons has been suggested [57], [58], [59], confirmatory mechanistic evidence was missing. Microglial cells are the primary immune cells of the central nervous system, highly similar to macrophages [60], and were originally thought to be a dormant immune population activated under pathological circumstances, like inflammation or infection [61]. However, microglia is also involved in physiological functions, such as mediating the communication between neuronal cells [62]. Building on the knowledge that Rank pathway is active in microglial cells [63], [64], [65], we now demonstrate that microglia of the hypothalamus regulates GnRH function through Rank signaling. Although scRNAseq analyses have effectively described neuronal subpopulations at the hypothalamic level, the characterization of microglial subpopulations remains limited due to microglia scarcity [20]. Our results now reveal that hypothalamic microglia exist in three distinct stages, with the most abundant being an activated subpopulation that is highly dependent on Rank signaling for its maintenance and functionality. Rank depletion not only reduces the relative abundance of the activated microglia subpopulation, but also compromises its functionality by impairing complement activation, protein translation and energy homeostasis. Alterations in protein synthesis disrupt microglia priming, phagocytosis, motility and synapse regulation [66], [67]. Moreover, Trem2/Tyrobp signaling pathway orchestrates microglial function, complement system and phagocytosis, pathways that are strongly affected upon Rank loss.

Rank loss not only causes profound transcriptomic changes in the microglia but also morphological alterations in the ME but not in ARC or PoA. The ME is the projection field of GnRH neurons, which plays a precise regulation of GnRH release. The amoeboid morphology caused by Rank loss in the ME was suggestive of hyperactive/inflamed microglia. Paradoxically, the reduction in microglia-GnRH contacts, GnRH engulfment by the microglia and the phagocytic marker CD68 was indicative of dysfunctional/hypoactive microglia with no signs of inflammation, in agreement with the transcriptomic results. Accumulating evidence indicates that microglia functionality cannot be inferred exclusively from its morphology and must be complemented by transcriptomic and functional analyses [68]. Our data provide evidence that engulfment of GnRH terminals by hypothalamic microglia in the ME is essential for their functionality and for the regulation of the HPG axis, in line with the known role of microglia in regulating synaptic communication and pruning of specific neurons [69].

In sum, our comprehensive studies in mouse models and human samples support that Rank-expressing microglia are key regulators of microglia activation, GnRH functionality and in turn, sexual development and fertility, uncovering RANK as a valuable therapeutic target in endocrine disorders and fertility syndromes, as well as a candidate gene for molecular diagnosis of CHH disorder.

## Acknowledgments

We thank all the patients who contributed to this study. We thank Vincent Prevot, Nathalie Journic, and the entire team of the Neuroendocrine Brain Development and Plasticity Laboratory (Inserm, University of Lille, France) for the discussions and support. We thank Robin Lovell-Badge (The Francis Crick Institute, UK), Andrea Vethencourt (Instituto Catalán de Oncología, Spain), Isabel Fariñas (Universidad de Valencia, Spain), María Casanova, Manuel Valiente, Rafael Fernandez-Leyro, Mercedes Robledo, Clara Reglero and Alberto Díaz (CNIO, Spain) for fruitful discussions. We thank IDIBELL and CNIO Animal Facilities for their assistance with mouse models, with a special remark to Miriam Garcia, Gema Luque and Isabel Blanco; the Genomics Unit, in particular Orlando Dominguez, and the Bioinformatics Unit, for their support with scRNAseq experiments. We thank Sagrario Ortega (CNIO) for the generation of Rank^e1-/-^ mouse model, Ana Semiao Rocha (IDIBELL) for supervising the generation of the animal models, Vivian Capilla-González (CNIO) for her help in the odor test experiments and Laura Álvaro, Sergi Velasco and all the members of the EGS laboratory for their valuable assistance during the development of the experiments and discussions of the results.

## Funding

Grants to EGS by European Research Council (ERC) under the European Union’s Horizon 2020 research and innovation programme (grant agreement no. 682935), Caixa Research Health 2023 Grants La Caixa Foundation (no. HR23-00361), Spanish Ministry of Science and Innovation, Agencia Estatal de Investigación (AEI) (SAF2017-86117-R, PID2020-116441GB-I00 and PID2023-152798OB-I00) co-funded by FEDER funds/ European Regional Development Fund (ERDF) (a way to build Europe) and PID2023-149244NB-I00 grant to MTS. FPU grant to A. Collado-Solé (FPU2018/00546) and FPI to A. Barranco (PRE2018-086522), Miguel Servet to CF (CP24_00036) Instituto de Salud Carlos III (ISCIII).

## Author contributions

Conceptualization: ACS, EGS

Methodology: ACS, NB, JZ, GSA, CF, FRP, CGL, YZ, VL, BMR, AB, GY

Investigation: ACS, NB, JZ, GSA, CF, FRP, GS, AJ, KR, JAS, APB, RFC, MTS, NP

Funding acquisition: EGS, MTS, NP, RFC, APB

Supervision: EGS, NP, MTS, RFC, APB

Writing – original draft: ACS, EGS

Writing – review & editing: all authors

## Competing interests

Authors declare that they have no competing interests.

## Data and materials availability

The scRNAseq results obtained in this work have been deposited at GEO, GSE240379.

## Supplementary Materials

Materials and Methods

Figs. S1 to S14

Supplementary Tables S1 to S7

## Notes

### Competing Interest Statement

The authors have declared no competing interest.

