## Supplementary material for "Microglia Rank signaling regulates GnRH function and the Hypothalamic-Pituitary-Gonadal axis": Suplemmentary information

Science
AAAS

### Supplementary Materials for

#### Microglia Rank signaling regulates GnRH function and the Hypothalamic-Pituitary-Gonadal axis

Alejandro Collado-Sole *et al.*

**The PDF file includes:**

Materials and Methods
Figs. S1 to S14
Tables S1 to S7
References

#### Materials and Methods

##### CHH Patients

The CHH study group consisted of 564 individuals (286 with Kallmann syndrome and 278 with normosomic CHH). The diagnosis of CHH was determined by three factors: i) absence or incomplete development of puberty by the age of 17; ii) low or normal levels of gonadotropins with low levels of testosterone or estradiol;; and iii) abnormal function of the anterior pituitary gland and abnormal imaging of the hypothalamic-pituitary region [1]. Olfaction was assessed through either self-reported information or formal testing [2]. Genetic testing was conducted on probands and their family members whenever available. This study was approved by the ethics committee of the University of Lausanne and all participants provided written consent prior to their involvement.

##### CHH Genetic analyses

We extracted genomic DNA from peripheral blood samples using the Puregene Blood Kit (Qiagen) as per the manufacturer's instructions. Exome capture was carried out using SureSelect All Exon capture v2 or v5 (Agilent Technologies), and the samples were sequenced on the HiSeq2500 (Illumina) at BGI (BGI, Shenzen). We used an in-house pipeline to analyze the raw sequences (FASTQ files) that utilized the Burrows-Wheeler Alignment algorithm (BWA) [3] for mapping the reads to the human reference sequence (GRCh38) and the Genome Analysis Toolkit (GATK) [4] to detect single nucleotide variants (SNVs) and insertion/deletions. The identified variants were annotated using Annovar version 20191024 [5] and gnomAD v4.0 for minor allele frequency (MAF), CADD [6] and AlphMissense [7] for pathogenicity scores. We selected variants passing multiple quality filters, with a coverage and quality score higher than 10 and 30 respectively, an allelic depth ratio higher than 0.2, and not present in segmentally duplicated regions [8]. Furthermore, rare variants present frequently in a local genetic database of healthy individuals (n=300), in excess of five pedigrees in the cohort, or flagged in gnomAD were considered systematic sequencing artifacts and discarded. We established a MAF threshold of 0.1% and excluded all variants with a higher popmax MAF in gnomAD. This is accounting for CHH prevalence in the general population [1] and recessive genes. For RSVs in RANK and RANK metagenes, we further applied pathogenicity filtering by retaining variants with either a CADD score > 20 or an AlphaMissense score > 0.5, in order to exclude likely benign variants. In addition, mutations in known CHH genes, as assessed by ACMG criteria, were investigated in our CHH cohort.

##### Rank$^{e1-/-}$ generation

The Rank$^{e1-/-}$ mouse model was generated by inserting a CreERT2-IRES-EGFP-pA construct, followed by an frt-flanked PGK-neo cassette, immediately downstream of the ATG translation initiation codon of the *Rank* gene. The targeting vector was electroporated into G4 mouse embryonic stem cells, and clones were selected in 200 µg/ml of G418. Homologous recombinant clones were identified by Southern blot analysis of genomic DNA digested with the restriction enzymes EcoRV (5' recombination) and KpnI (3' recombination) using 5' and 3' external probes amplified from genomic DNA. The primers used to amplify the 5'-1 probe (335 bp) by PCR were as follows: Forward: 5'-ATTGTGGGAGGGGTAACTGG-3' and Reverse: 5'-AAAAGAAAACAGAGCAGCGGG-3'. The primers used to amplify the 3'-2 probe (372 bp) were as follows: Forward: 5'-GCCATGAGTTCAACCCCTCA-3' and Reverse: 5'-

CACGAGAGGTCTGGCTTGTT-3'. The band sizes obtained for the 5' homologous recombination were 7.2 kb (KI) and 18.0 kb (WT), and for the 3' homologous recombination, 15.0 kb (WT) and 11.0 kb (KI). Chimeras were generated by microinjection into CD1 8-cell embryos. Germ line male chimeras were crossed with Tg.CAG-Flpe females to generate offspring in which the selection cassette was deleted. Deletion was confirmed by Southern blot analysis of genomic DNA digested from ear punch biopsies with DrdI and using a DNA fragment of 361 bp (probe 3'-1) as a probe. The probe 3'-1 was amplified by PCR from genomic DNA using the following primers: Forward: 5'-TTTTCAGGGTGGAGCATCCC-3' and Reverse: 5'-CACCATTGCCCTGACCTGAT-3'. The band sizes obtained were as follows: KI (deleted): 8.6 kb, KI (undeleted): 7.0 kb, and WT: 15.77 kb. Rank$^{e1+/-}$ mice were viable, fertile, and without overt abnormalities. In contrast, Rank$^{e1-/-}$ mice were infertile and exhibited developmental phenotypes similar to other Rank$^{-/-}$ mouse models[9].

**Mice**

All research involving animals was performed at the IDIBELL and CNIO Animal Facilities in compliance with protocols approved by the IDIBELL and CNIO Committees on Animal Care and following national and European Union reg.ulations. Rank flox/flox (Rank$^{fl/fl}$) mice were provided by Dr. Joseph Penninger [10] and crossed with K5-Cre (Terutine et al., 1997) to generate Rank$^{K5\Delta/\Delta}$, Csf1r-Cre [11] to generate Rank$^{Csf1r\Delta/\Delta}$, Gnrh1-Cre (MGI:3691288) to generate Rank$^{Gnrh1\Delta}$/Δ, Cx3cr1-Cre [12] to generate Rank$^{Cx3cr1\Delta/\Delta}$, LysMCre (MGI: 1934631) to generate Rank$^{LysM\Delta/\Delta}$ ,UbcCreERT2 to generate Rank$^{iUbc\Delta/\Delta}$, Cx3cr1-CreERT2 (MGI:6724385) to generate Rank$^{iCx3cr1\Delta/\Delta}$ and Tmem119-CreERT2 (MGI:6758066) to generate Rank$^{iTmem119\Delta/\Delta}$ mouse models. The resulting Rank$^{iCx3cr1\Delta/\Delta}$ mouse model was crossed with the reporter mouse line Rosa26tdtomato (MGI 155793) to label the recombinant cells. Rank$^{-/-}$ mice (MGI1314891) were previously characterized [9]. Rank$^{-/-}$, Rank$^{ex1-}$/-, and Rank$^{Csf1r\Delta/\Delta}$ mice display a strong osteopetrotic phenotype and absence of tooth eruption. Therefore, these mice were provided with a special liquid diet, which was changed daily, and maintained together with their littermates. Some Rank$^{Cx3cr1\Delta/\Delta}$ mice also show defective teeth; therefore, all mice were maintained on the same diet- Genotyping primers are indicated in Table S7.

***In vivo* treatment**

CreERT2 activation from Rank$^{iUbc\Delta/\Delta}$, Rank$^{iCx3cr1\Delta/\Delta}$ and Rank$^{iTmem119\Delta/\Delta}$ mice was achieved by tamoxifen (TAM) administration intraperitoneally (100 mg/kg). Three injections of TAM were administered every other day to mice at 4 weeks of age or in adulthood (10–12 weeks) in both experimental and control groups, and the animals were sacrificed 3 weeks after the last injection. For experiments of microglia depletion, AIN-76A rodent diet (Research Diets,333#D10001; control) and the same with CSF1R antagonist PLX3397334 (MedChemExpress, #D13050910; 600 ppm in chow) were used ad libitum from the corresponding stage (from 4 to 8 weeks).

**Intracranial injection**

Mice (12-15 weeks) were anesthetized by intraperitoneal injection of ketamine/xylazine cocktail (ketamine 15 mg/kg BW/xylazine 3 mg/kg BW) and placed in a stereotaxic frame. Four-hydroxytamoxifen (4-OHT) 10 mM was injected in the median eminence (ME) bilaterally using a 32-gauge needle connected to a 1-ml syringe (Neuro-Syringe, Hamilton) and was delivered at a rate of 0.1 μl/min for 7 min (1 μl/injection site) according to the following coordinates: −1.5 mm posterior to the bregma, ±0.2 mm lateral to midline, and −6 mm below the surface of the skull as previously reported [13].

**Odor test**
The odor test experiments were conducted in a plastic container from Kaiser+Kraft (40 cm×30cm×23cm). The odorants were administered using a cotton swab, which was impregnated with the relevant odorant and inserted through a small opening located 6 cm above the floor on one of the side walls. A 5-minute adaptation period was initiated, during which the swab was presented without any odorant (water), followed by six consecutive presentations of Odorant A (habituation phase) and six consecutive presentations of Odorant B (dishabituation phase), each lasting 1 minute [14]. Olive and sunflower oil were used as A and B odorants, respectively.

**Fertility assay**
A male was considered fertile if it had the capacity to impregnate a control female within a maximum period of 90 days, resulting in pregnancy and successful birth of pups. A female was considered fertile if she could be impregnated by a control male within the same period. A male or female was considered subfertile if more time was required to become pregnant or to impregnate compared to control mice.

**Pubertal onset and estrous cycle**
To assess pubertal maturation phenotypically, we monitored the day of balano-preputial separation in males and vaginal opening in females, both established external markers of puberty onset. In females, the day of first estrous cycles was determined through daily vaginal cytology beginning from the day of vaginal opening. Vaginal smears were collected and spread onto glass slides air-dried. The slides were then examined under a light microscope to identify the stage of the estrous cycle, based on the presence of cell types characteristic of each phase [15].

**Evaluation of gonadal maturation**
Gonadal maturation was assessed by morphometric analyses of H&E staining of a representative slide of testes and ovaries [16], [17]. Assessment of the presence of corpora lutea was conducted as index of ovulation. In the case of the testicular maturation, analyses of the area of circular seminiferous tubules were used as an indicator of spermatogenesis progression.

**Cholesterol analysis in serum**
Cholesterol levels were measured in mouse serum according to the manufacturer's instructions in Pentra C200 Clinical Chemistry Analyzer (Cholesterol CP ABX Pentra, A11A01634).

**Enzyme-linked immunosorbent assay (ELISA) of serum samples**
Testosterone (KGE010, R&D Systems) and estradiol (KGE014, R&D Systems) levels were measured in mouse serum according to the manufacturer's instructions.

**Whole mount staining**
For whole mount analyses, inguinal mammary glands were collected at the time specified, fixed and stained with carmine dye [18].

**Cleared fat pad transplantation assay**

Mammary epithelial cells were isolated from mouse mammary glands as previously described [19]. Cells isolated from mammary glands were diluted 1:1 with Matrigel Matrix (BD Biosciences, San Diego, CA, for a final volume of 20-30 µL and injected in cleared mammary fat pad of 3-4 weeks old C57BL/6 mice. After 8 weeks, the transplanted fat pads were whole-mounted and carmine stained.

**Pharmacological studies in response to GnRH and Kiss1 treatment**

For hormonal LH assays, blood samples were obtained from mouse tail at the designated times after intraperitoneal injection as reported in [20]. We analyzed time-course Lh response to intraperitoneal injection of GnRH (0.25 nmol/animal) and Kisspeptin-10 (7.5 nmol/animal). Animals were allowed to recover for 72 hours between tests. For each sample, 4 µL of whole blood was diluted 46 µL of 0.1 M phosphate-buffered saline (PBS) with 0.05% Tween 20, snap-frozen on dry ice, and stored at -80ºC. Mice were handled (5-10 min) every week before 3wk before blood sampling, to habituate them for tail-tip bleeding.

**Highly sensitive assay of LH levels in blood**

LH levels in blood were measured using a highly sensitive enzyme linked immune sorbent assay (ELISA) as reported in [21]. We use 50 µL of capture antibody diluted 1:1000 (Bovine LHβ 518B7 monoclonal Ab obtained from Lillian E Sibley @ UC Davis) and 50 µL of biotinylated detection antibody (Mouse Monoclonal LH antibody (Medix, 5303 SPRN-5) at 1:2000 in blocking Buffer 5% SMP-PBS-T (0.05% Tween-20). In addition, each well was incubated with 50 µL of Poly-HRP Streptavidin (Thermo Fisher, Cat# N200) and 100µl of OPD (o-Phenylenediamine) (1 Tablet-20 mg of OPD in 48ml of Citrate buffer +20µl of 30% $H_2O_2$; Sigma Aldrich).

**Microglia cell line and primary mouse microglia culture**

The microglial cell line BV2 (ABC-TC212S, Accgene) was grown in DMEM medium with 10% fetal bovine serum (Gibco) and 100 UI/ml penicillin and 100 µg/ml streptomycin (Gibco) in a water-saturated atmosphere of 5% $CO_2$ and 5% air. Microglia primary culture was obtained from a mixed astromicroglial primary culture from newborns C57BL/6 mice [22]. The day before the procedure, 6-well plates were coated with 10% of Poly-D-Lysine and incubated overnight at 37°C. Brains from neonatal mice (P0-P5, preferably P3) were carefully extracted under a dissection microscope, ensuring the removal of meninges to prevent contamination with perivascular macrophages. Brains were stored in F50 Hibernate solution until trypsinization. Brain tissues were mechanically dissociated using scissors and pipetting and chemically dissociated with trypsin for 7 minutes at 37°C in a thermomixer. Following digestion, the brain tissue was further homogenized by pipetting. The dissociated tissue was inactivated using Micro Full Medium (DMEM-F12 (1:1), 10% heat-inactivated FBS, 1% MEM NEAA, gentamicin (10 µg/ml), 100 UI/ml penicillin and 100 µg/ml streptomycin, filtered through a 40 µm nylon filter, and seeded into Poly-D-Lysine-treated wells (previously washed twice with water and left to dry). One day after seeding, 1 mL of fresh Micro Full Medium was added to the wells without washing, allowing cells to adhere. Media changes were performed every 2–4 days without washing the cultures. Approximately 20 days after seeding, a gentle trypsinization (0.25% trypsin with 1 mM EDTA) was performed to remove astrocytes from the culture. The cultures were incubated at 37°C for 20–45 minutes, until separation occurred. Trypsin was then inactivated with Micro Full Medium. Subsequently, astrocyte-conditioned medium (collected from the astro-microglial plates and diluted 1:1 with

Micro Full) was added, and cultures were incubated for 24 hours. Then, cells were treated with Rankl (100 ng/mL) for 6 hours. The cell pellet was then collected for RNA extraction.

**Tissue histology and immunostaining in paraffin blocks**

Mouse tissue samples (mammary gland, bone, testes and ovaries) were fixed in formalin overnight at 4ºC and embedded in paraffin according to the CNIO Histopathology Unit protocols. Bone samples were decalcified by the CNIO Histopathology Unit prior to embedding. Five-µm sections were cut for histological analyses processed for H&E staining or immunostaining. Monoclonal anti-ERα antibody (non-diluted medium from hybridoma) was generated by CNIO Monoclonal Antibodies Unit and immunohistochemistry of mammary gland was performed by CNIO Histopathology Unit.

**Brain tissue histology**

Adult mice were anesthetized with 50–100 mg/kg of Ketamine-HCl and 5–10 mg/kg Xylazine-HCl and perfused transcardially with 20 mL of PBS, followed by 50–100 mL of 4% PFA (pH 7.4). Brains were collected and fixed in the same fixative for 24 hours at 4°C, cryoprotected in 30% sucrose, embedded in optical cutting temperature (OCT) embedding medium (Tissue-Tek), frozen on dry ice, and stored at −80°C until use. Tissues were cryosectioned (Leica cryostat) between 45–80 µm. Coronal sections were washed in PBS and incubated for 60 minutes in blocking solution (0.3% BSA + 0.3% Triton X-100 in 1× PBS for PoA, or 0.3% Triton X-100, 10% normal donkey serum, and 1% BSA for mediobasal hypothalamus). Sections were then incubated overnight with primary antibodies diluted in the respective blocking solution: rabbit anti-GnRH (1:1000, 269501-AP, Proteintech), goat anti-Iba1 (1:500, AB5076, Abcam), chicken anti-Iba1 (1:1000, 234009, Synaptic Systems), and rat anti-CD68 (1:500, MCA1957GA, Bio-Rad). TdTomato fluorescence was detected by endogenous signal. After primary antibody incubation, sections were rinsed three times in PBS and incubated for 120 minutes at room temperature with fluorochrome-conjugated secondary antibodies (1:1000, Jackson ImmunoResearch). Finally, sections were washed, mounted with Permount containing DAPI, and coverslippe. For quantification of GnRH neuron population in the PoA, images were acquired using a Zeiss Axio Imager Z2 ApoTome microscope (Zeiss, Germany), as previously reported [23]. The quantification was performed on one-third of the mouse brain and then normalized to account for the total GnRH population To analyze the number and morphology of microglia in the medio basal hypothalamus, 3D confocal images were acquired using the z-stack function on an LSM 700 confocal microscope with a 40× oil-immersion objective (Zeiss, Oberkochen, Germany), a Leica Stellaris 8 microscope with 20×/63× glycerol-immersion objectives, or a Leica TCS SP8 confocal microscope with a 40× oil-immersion objective. Maximal Z-projections were generated using the same number of planes for each staining. Three-dimensional reconstruction was performed with Bitplane Imaris 9 software (Bitplane, Zurich, Switzerland) and Imaris x64 9.6.0 image analysis software (Oxford Instruments, Concord, MA). Images were first subjected to background subtraction and then processed using the surface and filament modules to reconstruct microglia, microglial cell bodies, microglial processes, GnRH, and CD68. Although most of the analysis was automated by the software, the origin of each process was independently verified to ensure correct cell assignment, and any erroneous connections were manually removed.

**iDISCO (immunolabeling-enabled three-dimensional imaging of solvent-cleared organs)**

Briefly, tissues were fixed in 4% PFA by perfusion and stored in PBS. Samples were dehydrated in a methanol/PBS gradient (20%, 40%, 60%, 80%, and 100% twice, 1 hour each) and then delipidated with dichloromethane (2:1 dichloromethane/methanol) overnight at 4 °C. The following day, samples were bleached in a hydrogen peroxide/methanol solution (1:6 $H_2O_2$ to 5:6 methanol) overnight to reduce autofluorescence. On the third day, the samples were hydrated in a methanol/PBS gradient (100%, 80%, 60%, 40%, 20%, PBS; 1 hour each). After rehydration, samples were permeabilized and blocked in PBSGT (PBS, 0.2% gelatin, 1% Triton X-100, 0.05% sodium azide) for 4 days and then incubated with GnRH antibody (Proteintech, 26950-1-AP) for 14 days. Subsequently, samples were washed six times for 1 hour each in PBS + 1% Triton X-100 and then incubated with Alexa Fluor 555 anti-rabbit secondary antibody (Life technologies, SA) in PBSGT for 1 week. After secondary antibody incubation, samples were washed six times for 1 hour each in PBS + 1% Triton X-100. All immunolabeling steps were performed with gentle rotation and protection from light. For tissue clearing, samples were dehydrated again in a methanol/PBS gradient (20%, 40%, 60%, 80%, and 100% twice, 1 hour each) and then delipidated with dichloromethane (2:1 dichloromethane/methanol) overnight at 4 °C. The following day, samples were immersed in dibenzyl ether (DBE) to achieve optical transparency. Cleared tissues were stored in fresh DBE at room temperature, protected from light, and imaged as whole mounts. Three-dimensional imaging was obtained as reported[23]. Imaging was performed using an Ultramicroscope I (LaVision BioTec) controlled with ImspectorPro software (LaVision BioTec). The light sheet was created with a laser in combination with two cylindrical lenses. A binocular stereomicroscope (MXV10, Olympus) equipped with a 2× objective (MVPLAPO, Olympus) allowed imaging at multiple magnifications (1.6×, 4×, 5×, and 6.3×). Samples were placed in a 100% quartz imaging chamber (LaVision BioTec) filled with DBE and illuminated laterally using the full width of the laser sheet. Image acquisition was performed with a PCO Edge SCMOS CCD camera (2,560 × 2,160 pixels, LaVision BioTec) with a z-step size of 2 μm. The resulting TIFF image stacks were converted to Imaris-compatible files (Imaris FileConverter, Bitplane) for three-dimensional reconstruction and imported into Imaris (Bitplane) for visualization and generation of snapshots.

**RNA isolation, RT-PCR and gene expression analysis**
Total RNA was isolated from mouse tissue using Maxwell® RSC simplyRNA Tissue Kit (AS1340 Promega). One thousand ng (testis/ovary/cell lines) or 500 ng (pituitary gland/hypothalamus) of RNA was reverse-transcribed into cDNA using 200 U Superscript II plus random hexamer oligos (Invitrogen). Different genes were amplified with LightCycler® 480 Probes Master (Roche, 04707494001) and a LightCycler® 480 thermocycler (Roche) and normalized relative to the *Hprt1* or *Gapdh* mouse keeper gene. Primers sequences are indicated in Table S7.

**Public scRNAseq analyses**
Analyses of gene expression in mouse hypothalamus [24], were conducted using the interactive CellxGene viewer (https://www.mrl.ims.cam.ac.uk), while analyses of gene expression in human hypothalamus [25].were performed using R/Seurat.

**Metagene analyses and correlations**
Metagenes and correlation analyses provided in the manuscript were generated by GEPIA2 [26]. Pearson's correlation coefficient was used as the statistical test for the correlation analyses. The microglial gene signature was extracted from PanglaoDB[27], a publicly available single-cell

RNA sequencing database that provides well-characterized, cell-type specific gene signatures and is widely recommended for cell and tissue identification in scRNA-seq studies.

**scRNAseq of mouse hypothalamus**

Single-cell suspensions were prepared from the hypothalamus of $Rank^{iUbc\Delta/\Delta}$ mice enriched in microglia. These mice were treated with tamoxifen (100 mg/kg) at puberty onset and analyzed four weeks after the first treatment. The weight of the testes was measured as an indicator of efficient Rank depletion. Hypothalamic tissue from 5 control and 5 $Rank^{iUbc\Delta/\Delta}$ mice was pooled on ice. The tissue was then mechanically and enzymatically digested using DMEM/F12 (Gibco), 0.3% collagenase A (Sigma), 2.5 U/ml dispase (Sigma), 2 mM HEPES, and Penicillin/streptomycin (ThermoFisher Scientific) for 30 minutes at 37 ºC, with the digestion buffer changed every 10 minutes. After digestion, the cells were filtered through a 70 µm filter, and centrifuged at 300 g for 5 minutes at 4 ºC. Erythrocytes were removed using ACK buffer, followed by using 40% Percoll to eliminate myelin debris. Subsequently, single cells from the hypothalamus were sorted to remove debris and select live cells for further analysis. Cell sample was loaded onto a 10x Chromium Single Cell controller chip B (10x Genomics) as described in the manufacturer's protocol (PN-1000121, Chromium Next GEM Single Cell 3' GEM, Library & Gel Bead Kit v3.1). Libraries were sequenced on the Illumina NextSeq 550 platform (with v2.5 reagent kits) with paired-end sequencing (28 bp + 56 bp bases).

**Data processing, normalization, and clustering annotation**

Bollito [28] pipeline was employed to perform initial steps of the analysis as follows: sequencing quality was checked with FastQC (http://www.bioinformatics.babraham.ac.uk/projects/fastqc/); reads were aligned to the mouse reference genome (GRC39m from GENCODE [29]) with STARsolo (STAR v2.7.1) [30]; Seurat v3.2.3 [31] software was used to check the quality of sequenced cells, and perform data normalization, dimensionality reduction and clustering. Cells expressing less than 200 features and more than 2x the median of features were filtered out. Moreover, a maximum cutoff of 10% and 40% were set for mitochondrial and ribosomal percentage, respectively. Mitochondrial percentage and merge effect were regressed out in order to minimize their effect on the samples. To perform the dimensionality reduction, the top 2500 variable genes are considered, and PCs and number of neighbors are both set to 20. The microglial subpopulation was subsetted for further clustering using the same approach. Annotation of the resulting clusters was first performed using singleR v2.0.0 [32] according to data provided by previous public brain datasets [33], [34] and afterwards this annotation was manually curated based on marker genes extracted from recent literature regarding hypothalamic cell subpopulations [24], [35].

**Differential gene expression analysis.**

Differential gene expression analysis (DEA) was performed using Seurat through the default Wilcoxon test, and significance is defined by a 0.05 adjusted P value threshold and an absolute log2-fold change (log2FC) > 0.25. In addition, default minimal threshold of 10% of expression in (at least) one of the compared populations is conserved.

**Gene Set Enrichment Analysis**

Gene Set Enrichment Analysis (GSEA) was used to interpret gene expression data through GSEAPreranked [36] on a preranked gene list sorted according to log2FC resulting from previous

DEA. Those gene sets with significant enrichment levels (FDR q-value < 0.25 or 0.05) were considered. 1.000 permutations are performed and a minimal and maximum number of genes per reference were defined as 10 and 500 respectively.

**Statistical analysis and reproducibility**

Statistical analyses were performed using GraphPad Prism software version 8. Data are represented as the mean ± S.E.M. The measurements were taken from different mice, avoiding technical replicates. When comparisons were made between two experimental groups, an unpaired, two-tailed Student's t-test was used. When comparing multiple variables between two experimental groups, an analysis of variance (ANOVA) was employed, followed by post hoc tests for multiple comparisons (Tukey). For the analysis of pubertal onset, a Gehan–Breslow–Wilcoxon matched-pairs test was applied
